# MicroRNA-27a-5p inhibits proliferation, migration and invasion and promotes apoptosis of Wilms tumor cell by targeting PBOV1

**DOI:** 10.1101/2021.08.25.457737

**Authors:** Zheng-Tuan Guo, Qiang Yu, Chunlin Miao, Wenan Ge, Peng Li

## Abstract

Wilms tumor is the most common type of renal tumor in children. MicroRNAs (miRNA) are small non-coding RNAs that play crucial regulatory roles in tumorigenesis. We aimed to study the expression profile and function of miR-27a-5p in Wilms tumor. MiR-27a-5p expression was downregulated in human Wilms tumor tissues. Functionally, overexpression of miR-27a-5p promoted cell apoptosis of Wilms tumor cells. Furthermore, upregulated miR-27a-5p delayed xenograft Wilms tumor tumorigenesis in vivo. Bioinformatics analysis predicted miR-27-5p directly targeted to the 3’-untranslated region (UTR) of PBOV1 and luciferase reporter assay confirmed the interaction between miR-27a-5p and PBOV1. The function of PBOV1 in Wilms tumor was evaluated in vitro and knockdown of PBOV1 dampened cell migration. In addition, overexpression of PBOV1 antagonized the tumor-suppressive effect of miR-27a-5p in Wilms tumor cells. Collectively, our findings reveal the regulatory axis of miR-27-5p/PBOV1 in Wilms tumor and miR-27a-5p might serve as a novel therapeutic target in Wilms tumor.

## Introduction

Wilms tumor is the commonest renal carcinoma mostly happened in children under age 5(Rivera and Haber, 2005). The survival of Wilms tumor patients has been significantly improved from less than 30% to more than 90% in the past decade due to modern therapeutic strategies and technology(Szychot et al., 2014; Lopes and Lorenzo, 2017). However, the current therapies such as radiotherapy and chemotherapy have serious side effects with poor efficacy in patients with tumor metastasis(Pritchard-Jones, 2002; Ehrlich et al., 2009; Akakin et al., 2016). Meanwhile, large-scale next-generation sequencing has identified multiple mutations of candidate driver in Wilms tumor(Gadd et al., 2017). Thus, it is important to further understanding the molecular mechanism of Wilms tumor oncogenesis and metastasis and develop new treatment strategies.

MicroRNAs (MiRNAs) are small non-coding RNAs that play crucial regulatory roles in various biological processes including tumorigenesis(Peng and Croce, 2016; O’Brien et al., 2018). MiRNAs post-transcriptionally regulate their target gene expression via binding to the 3’-UTR of target mRNAs(O’Brien et al., 2018). The expression and function of miRNAs in Wilms tumor have also been investigated(Yu et al., 2016). MiRNA microarray profiling results from 36 Wilms tumor of different subtypes and normal kidney tissues demonstrated that various miRNAs were dysregulated in blastema Wilms tumor and regressive subtype of Wilms tumor(Ludwig et al., 2016). Those miRNAs function as oncogenes, tumor suppressors, or mediate the chemo-sensitivity in Wilms tumor(Ludwig et al., 2016). Watson JA et al. also reported that miRNAs could predict the chemo-responsiveness in Wilms tumor blastema, with 29 miRNAs identified to be markedly differentially expressed in post-treatment high-risk and intermediate-risk patients(Watson et al., 2013). While miR-483-3p functions as an oncogene and promotes the development and chemo-resistance of Wilms tumor, miR-27 was reported to be downregulated in Wilms tumor(Watson et al., 2013; Wegert et al., 2015). However, the detailed functional role and molecular mechanisms of miR-27 in Wilms tumor are not fully understood.

In this study, we evaluated the expression profile and function of miR-27-5p in Wilms tumor and cell lines. The results demonstrated that miR-27a-5p was low-expressed in human Wilms tumor tissues and cells. Overexpression of miR-27a-5p inhibited cell proliferation, cell migration and invasion and promoted cell apoptosis. Moreover, our data revealed that miR-27a-5p suppressed tumorigenesis via negatively regulating PBOV1. In summary, our findings suggest that miR-27a-5p might serve as a novel therapeutic target in Wilms tumor.

## Materials and Methods

### Patient specimens

Twenty pairs of Wilms tumor and adjacent normal kidney tissues were obtained from patients who underwent surgery at the Second Affiliated Hospital of Xi’an Jiaotong University (Xibei Hospital) between March 2019 and September 2019. All Wilms tumor tissues were histopathologically confirmed and classified based on the American National Wilms Tumor Study 5 typing and TNM staging system. Tissues were snap-frozen and stored in liquid nitrogen until further analysis. The study was reviewed and approved by the Ethics Committee of Xibei Hospital and all patients provided the written informed consent.

### Cell culture

Wilms tumor-derived renal cancer cell line WiT49, STA-WT3ab, RM1, PSU-SK-1 and control HEK 293T cells were purchased from American Type Culture Collection (ATCC, USA). Cells were maintained in DMEM medium supplemented with 10% fetal bovine serum (Gibco, USA) and 1% penicillin/streptomycin (Gibco, USA) at 37 °C in a 5% CO_2_ incubator.

### Transfection

Transfection was conducted using Lipofectamine 2000 (Invitrogen, USA). MiR-27a-5p mimic or negative control (NC), miR-27a-5p inhibitor and NC were obtained from GenePharma (Shanghai, China). PBOV1 siRNA and scramble control were purchased from Rubibio Company (Guangzhou, China). pcDNA 3.1-PBOV1 was obtained from Genecopoeia (Maryland, USA).

### Lentivirus infection and generation of stable cell line

To generate stable cell line overexpressing miR-27a-5p or miR-NC, WiT49 cells were infected with lentivirus-miR-27a-5p mimic or lentivirus-miR-NC lentivirus particles. Lentivirus-miR-NC vector or lentivirus-miR-27a-5p mimic vector, together with the helper plasmids pHelper 1.0 (Gag and Pol) and pHelper 2.0 (VSVG) were transiently cotransfected into HEK 293T cells using Lipofectamine 2000 (Invitrogen, USA) to package lentivirus particles, respectively. When WiT49 cells were grown in the logarithmic growth phase, cells were infected with lentivirus particles with a multiplicity of infection of 70. Then, limited dilution method was used and cells were cultured with puromycin (2 µg/mL) and screened for 14 days to select the stable cell lines overexpressing miR-27a-5p or miR-NC.

### RT-qPCR

To analyze the relative expression level of miR-27a-5p and PBOV1, RNA was purified from Wilms tumor tissues or cultured cells using a miRNeasy Kit (Qiagen, German) and then reverse transcribed into cDNA using MiR-X miRNA Synthesis kit (Clontech, USA) or SuperTranscriptase III (Invitrogen, USA) following the manufacturers’ instructions. Quantitative PCR was conducted using the SYBR Green mix (Takara, Japan) on a Bio-Rad Real-time CFX96 system. U6 snRNA and GAPDH were used as the internal control. The gene expression was calculated using the 2^-ΔΔCt^ method. The following primers were used: hsa-miR-27a-5p: 5’-AGGGCTTAGCTGCTTGTGAGC-3’; hsa-PBOV1-Forward: 5’-AGTTCGAGACCAGCCTGACCAG-3’, hsa-PBOV1-Reverse: 5’-TTCAAGCAATTCTCCGCCTCAGC-3’; hsa-SPARC-Forward: 5’-CAAGAAGCCCTGCCTGATGAGAC-3’, hsa-SPARC-Reverse: 5’-TTCCGCCACCACCTCCTCTTC-3’; hsa-GSDMA-Forward: 5’-ACGCTGCTGGATGTGCTTGAG-3’, hsa-GSDMA-Reverse: 5’-AGAGCCCTGCCGTTCCCTTC-3’; hsa-ASB15-Forward: 5’-GCCTGGACATTAGGTTTGACC-3’, hsa-ASB15-Reverse: 5’-GTCAGGGGAGCCAGATAAGC-3’; hsa-UBXN4-Forward: 5’-CCTTCTGATGCTCCTCTAGAAG-3’, hsa-UBXN4-Reverse: 5’-GGAAACATGGTTGCTAACGAAA-3’.

### Western blot

Protein was prepared from tumor tissues or cultured cells using RIPA buffer (Beyotime, China) and quantified with a BCA protein assay kit (Pierce, USA). Western blot was performed using a standard protocol with primary antibodies against PBOV1 (Abcam, ab216045), β-actin (Abcam, ab8227). β-actin was used as a loading control.

### Luciferase reporter assay

Luciferase reporter vectors containing WT or mutated 3’-UTR of PBOV1 were constructed based on the backbone vector pGL3-Luc. WiT49 cells were co-transfected with reporter vectors, and miR-NC or miR-27a-5p mimic. After 48 hours, luciferase activity was measured with a Dual-luciferase reporter assay kit (Promega, USA).

### CCK-8 assay

Transfected WiT49 or STA-WT3ab cells were cultured in 96-well plates and 10 μl Cell Counting Kit-8 (CCK-8, Dojindo, Japan) reagent was added and cultured for 2 more hours. The absorbance (450 nm) was recorded 2 hours later.

### Cell apoptosis assay

Transfected WiT49 or STA-WT3ab cells were collected and stained with a BD apoptosis analysis kit (BD Bioscience, USA). After staining, cells were analyzed using the Cytoflex flow cytometry machine (Beckman Coulter), and the Annexin V+ Propidium iodide – cells were defined as apoptotic cells. The results were analyzed by Flowjo (Treestar, USA).

### Transwell assay

Transfected WiT49 or STA-WT3ab cells were resuspended in serum-free medium and seeded to the top chamber (Corning, USA) with or without pre-coating of Matrigel (BD Bioscience, USA). Complete medium with 10% FBS was added to the bottom chamber. After culture for 24 hours, the invaded or migrated cells were stained with 0.1% crystal violet (Solarbio, China) and counted.

### Xenograft tumor model

Ten male BALB/c nude mice (5-6 weeks old) were purchased from SLAC animal center (Shanghai, China) and randomly divided into 2 groups. WiT49 cells with stable overexpressing miR-27a-5p mimic or miR-NC were inoculated subcutaneously into nude mice, respectively. Tumor growth was recorded at indicated time points and calculated: Volume = length × width^2^/2. Mice were sacrificed and analyzed on day 22. The experiments were approved by the Animal Care Committee of Xibei Hospital.

### Statistical analysis

**Results** were shown as mean ± standard deviation (SD) from three independent experiments and analyzed by using GraphPad Prism V7.0 (Prism, USA). Student *t-*test and one-way analysis of variance (ANOVA) were conducted where necessary. A *p* < 0.05 was defined as statistically significant.

## Results

### MiR-27a-5p is downregulated in human Wilms tumor tissues and cells

To determine the expression of miR-27a-5p in Wilms tumor, we performed qPCR to examine the miR-27a-5p expression in twenty pairs of Wilms tumor tissues and adjacent control tissues (**Fig 1A**). MiR-27a-5p expression was significantly decreased in human Wilms tumor tissues (**Fig 1A**). Consistently, MiR-27a-5p expression was measured in different Wilms tumor cell lines and the results showed that Wilms tumor cell lines (WiT49 and STA-WT3ab) had markedly lower levels of miR-27a-5p than that in control cell line HEK 293T (**Fig 1B**).

**Figure 1.**
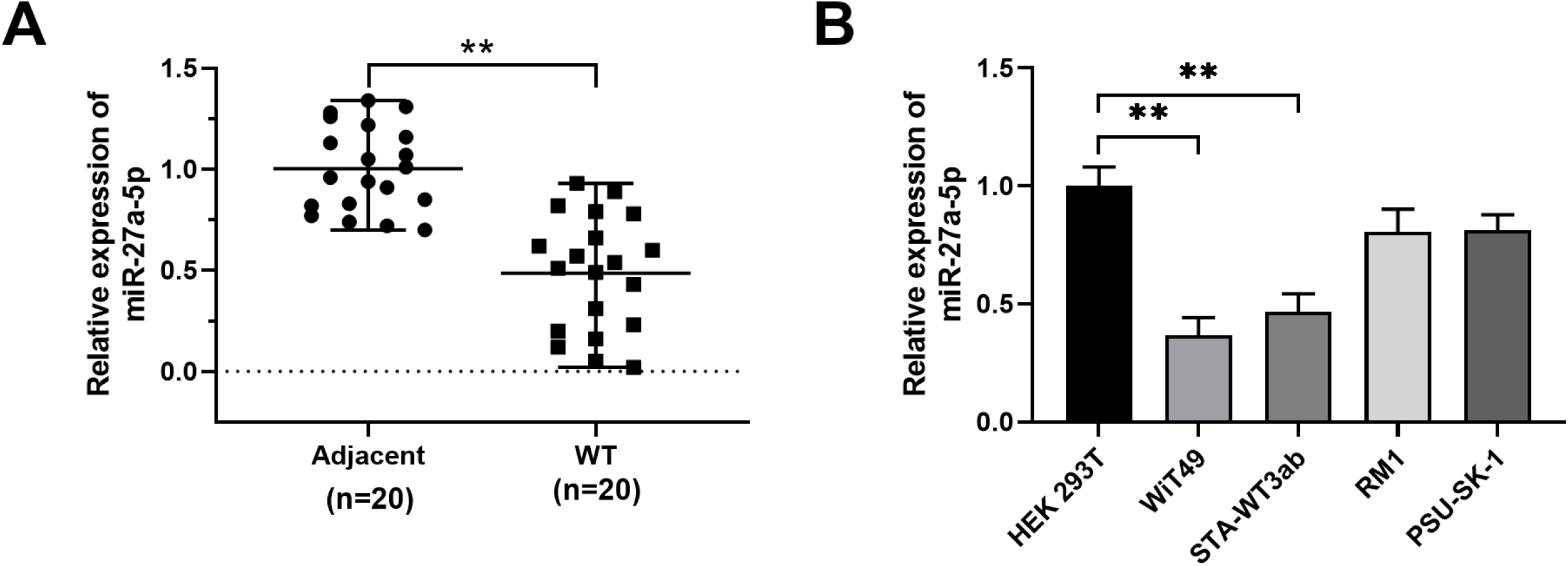
MiR-27a-5p expression in Wilms tumor tissues and cell lines. (A) Relative expression of miR-27a-5p was determined in twenty pairs of Wilms tumor tissues and adjacent normal control tissues by qPCR. (B) Relative expression of miR-27a-5p was determined in Wilms tumor cell lines (WiT49, STA-WT3ab, RM1, and PSU-SK-1) and control cell line HEK 293T by qPCR. ** *p* < 0.01.

### MiR-27a-5p inhibits proliferation, migration and invasion and promotes apoptosis in Wilms tumor cells

Then we performed functional assays to assess the role of miR-27a-5p in Wilms tumor cells. WiT49 and STA-WT3ab cells, which had relatively lower expression of miR-27-5p, were transfected with miR-27a-5p mimic to overexpress miR-27-5p (**Fig 2A**). Overexpression of miR-27a-5p remarkably repressed cell proliferation (**Fig 2B**) and promoted cell apoptosis (**Fig 2C**) in WiT49 and STA-WT3ab cells. Furthermore, compared with the negative control, miR-27a-5p mimic transfection significantly suppressed cell migration and invasion (**Fig 2D** and **2E**). These findings suggest that miR-27a-5p negatively regulates Wilms tumor development.

**Figure 2.**
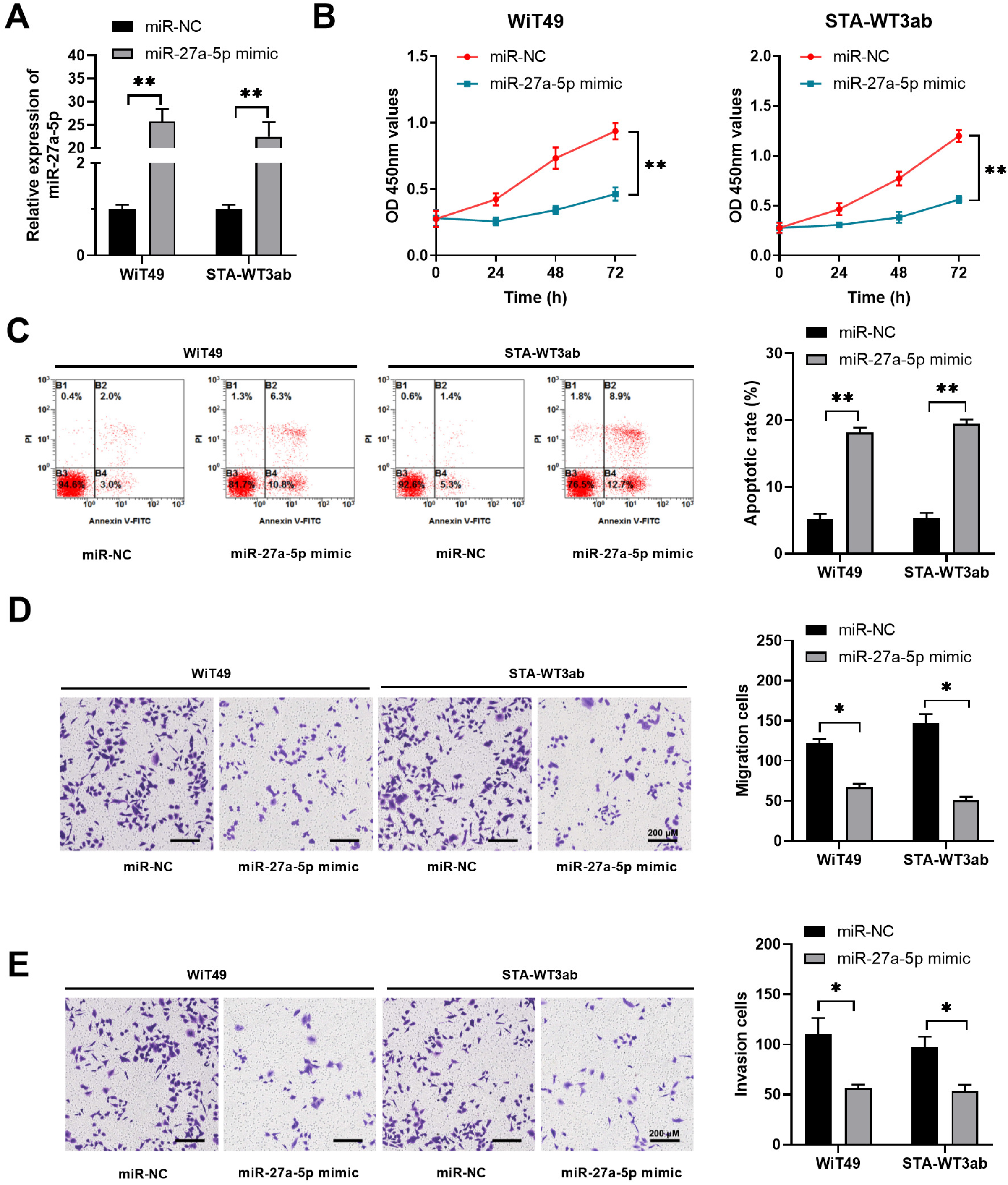
MiR-27a-5p inhibits proliferation, migration and invasion and promotes apoptosis in Wilms tumor cells. WiT49 or STA-WT3ab cells were transfected with miRNA negative control (miR-NC) or miR-27a-5p mimic. (A) The relative expression of miR-27a-5p in WiT49 or STA-WT3ab cells was analyzed by qPCR. (B) Cell proliferation was analyzed at indicated time points by CCK8 assay. (C) Cell apoptosis was analyzed by Annexin V/PI staining and flow cytometry. (D, E) Cell migration and invasion were assessed by transwell assay. * *p* < 0.05, ** *p* < 0.01 vs. miR-NC.

### MiR-27a-5p inhibits oncogenesis and metastasis of Wilms tumor in vivo

To verify the tumor suppressor role of miR-27-5p, in vivo xenograft model was established in nude mice by subcutaneous injection of WiT49 cells transfected with miR-27a-5p mimic or control miRNA. The results demonstrated the tumor inhibitory function of miR-27a-5p in vivo. MiR-27a-5p overexpression significantly inhibited Wilms tumor development (**Fig 3A**). Tumors developed from the miR-27a-5p mimic group showed a much smaller size, with a lower tumor weight in comparison with those developed from control cells (**Fig 3B** and **3C**). The upregulated miR-27a-5p expression was confirmed in tumors from the miR-27a-5p mimic group (**Fig 3D**).

**Figure 3.**
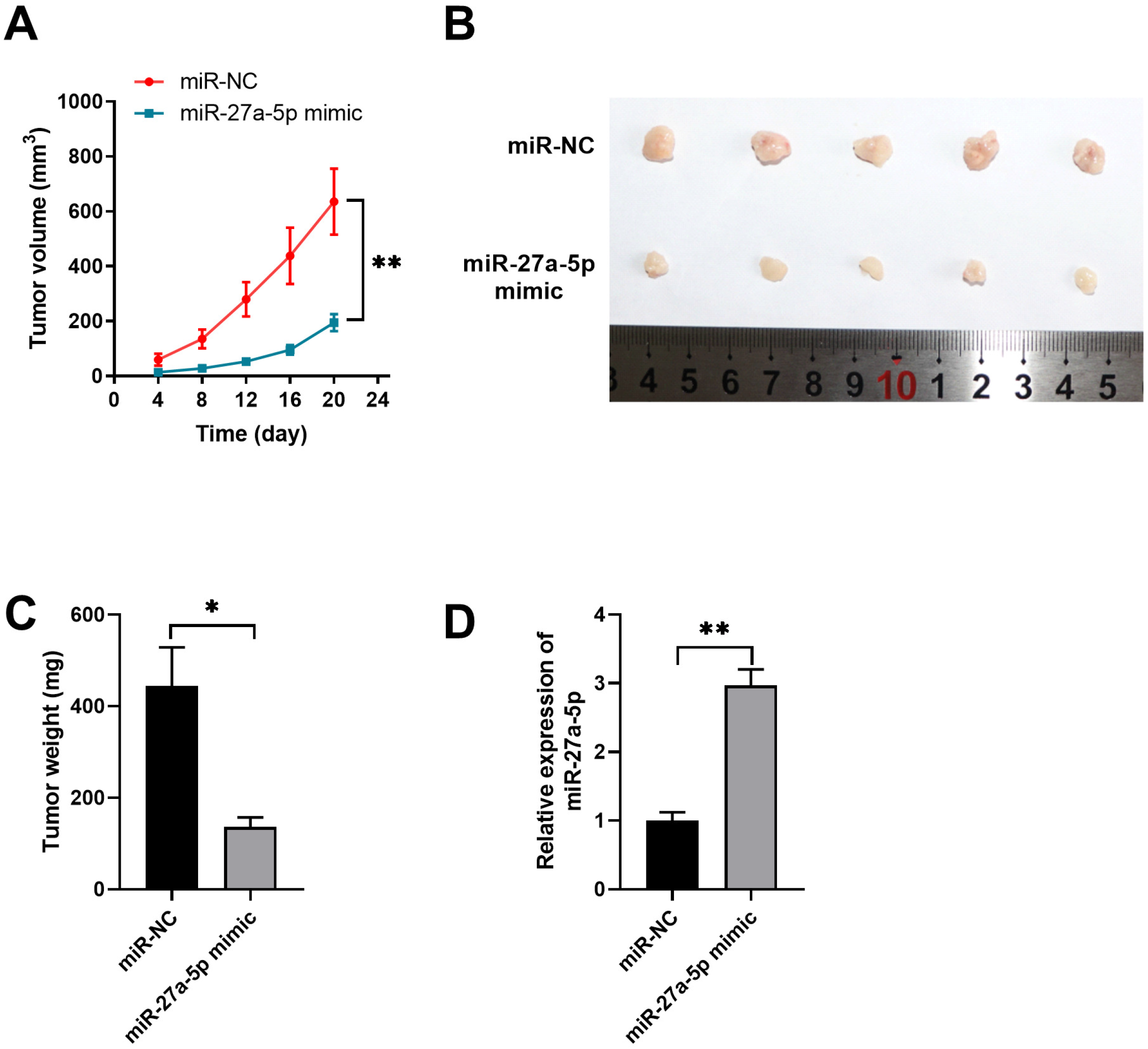
MiR-27a-5p inhibits Wilms xenograft tumor growth in vivo. WiT49 cells were transfected with negative control miRNA (miR-NC) or miR-27a-5p mimic and then inoculated into BALB/c nude mice to develop the xenograft Wilms tumor. (A) Tumor growth was monitored and measured at indicated time points. (B) Tumors were extracted and recorded on Day 22. (C) Tumor weights of xenograft were analyzed. (D) The relative expression of miR-27a-5p in xenograft tumors was analyzed by qPCR. * *p* < 0.05, ** *p* < 0.01 vs. miR-NC.

### PBOV1 is a direct target of miR-27a-5p in Wilms tumor cells

To explore the potential targets regulated by miR-27a-5p, we performed bioinformatics analysis using different online databases (TargetScan, miRDB, and miRWalk). As shown in **Fig 4A**, five genes including PBOV1, SPARC, ASB15, UBXN4, and GSDMA were predicted to the potential targets of miR-27a-5p. WiT49 cells were transfected with miR-NC or miR-27a-5p mimic and the relative expression of these potential targets was analyzed. PBOV1 was markedly downregulated by miR-27a-5p mimic (**Fig 4B**). In addition, miR-27a-5p had the putative binding sequences against 3’-UTR of the PBOV1 gene (**Fig 4C**). Luciferase reporter assay further validated the interaction between miR-27a-5p and WT 3’-UTR of PBOV1, as miR-27a-5p markedly inhibited the luciferase activity of reporter vector containing WT 3’-UTR of PBOV1 (**Fig 4D**). Moreover, we demonstrated that miR-27a-5p mimics significantly inhibited the mRNA and protein levels of PBOV1 in WiT49 or STA-WT3ab cells (**Fig 4E** and **4G**). In the contrast, inhibition of miR-27a-5p enhanced the expression of PBOV1 in WiT49 or STA-WT3ab cells (**Fig 4F** and **4H**).

**Figure 4.**
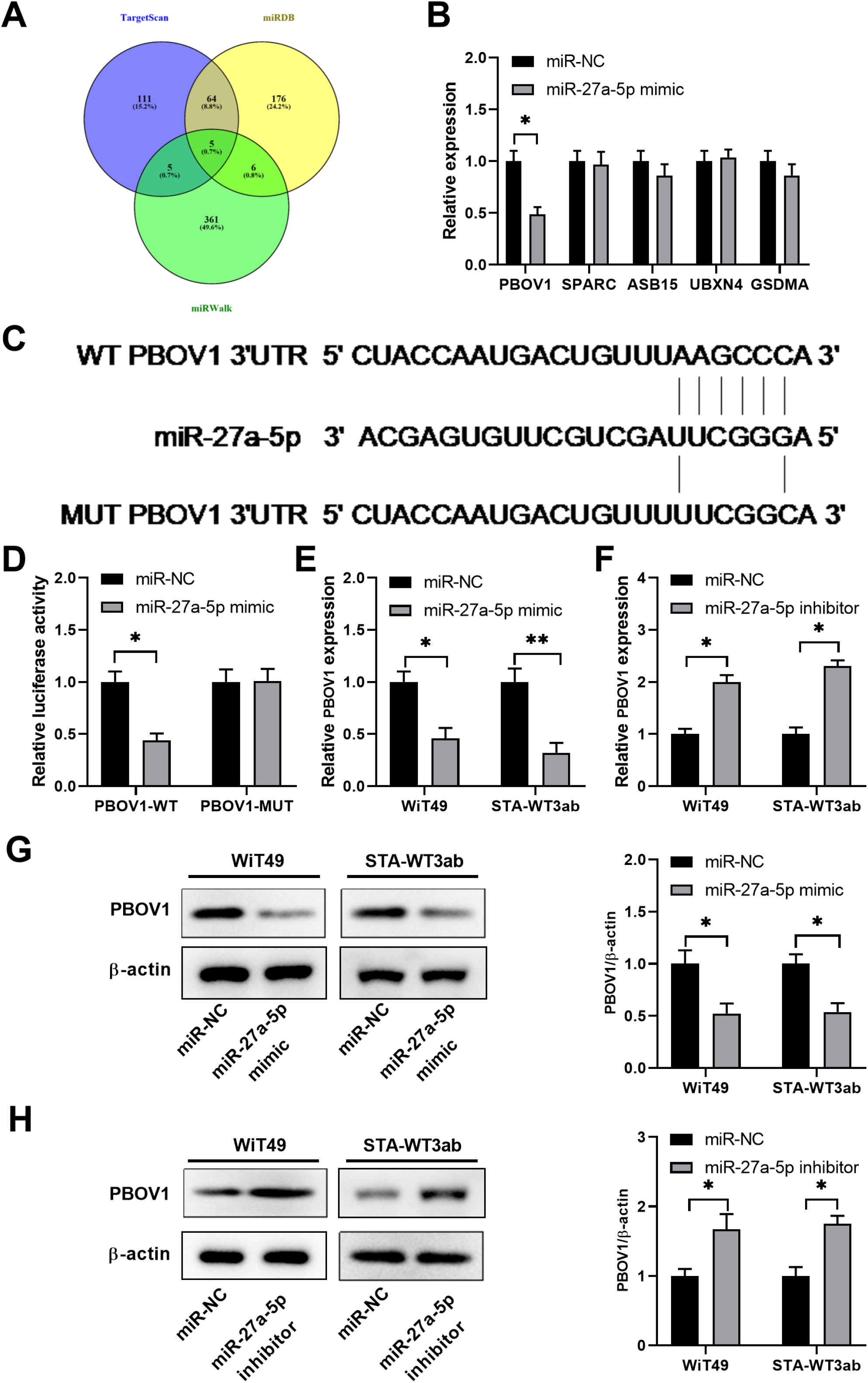
PBOV1 is a direct target of miR-27a-5p in Wilms tumor cells. (A) Bioinformatics analysis was performed to predict the potential targets of miR-27a-5p using online databases TargetScan, miRDB and miRWalk. (B) WiT49 cells were transfected with miR-NC or miR-27a-5p mimic. The relative expression of PBOV1, SPARC, ASB15, UBXN4 and GSDMA was analyzed by qPCR 48 hours later. (C) The predicted binding sequences between miR-27a-5p and WT or mutated 3’-UTR of PBOV1. (D) WiT49 cells were transfected with miR-NC or miR-27a-5p, together with luciferase reporter vectors containing WT or mutated 3’-UTR of PBOV1. The relative luciferase activity was analyzed 48 hours later. (E) WiT49 or STA-WT3ab cells were transfected with miR-NC or miR-27a-5p mimic. The relative expression of PBOV1 mRNA was analyzed by qPCR 48 hours later. (F) WiT49 or STA-WT3ab cells were transfected with miR-NC or miR-27a-5p inhibitor. The relative expression of PBOV1 mRNA was analyzed by qPCR 48 hours later. (G) WiT49 or STA-WT3ab cells were transfected with miR-NC or miR-27a-5p mimic. The relative expression of PBOV1 protein was analyzed by western blot 48 hours later. (H) WiT49 or STA-WT3ab cells were transfected with miR-NC or miR-27a-5p inhibitor. The relative expression of PBOV1 protein was analyzed by western blot 48 hours later. * *p* < 0.05, ** *p* < 0.01 vs. miR-NC.

### Knockdown of PBOV1 suppresses cell migration and invasion and promotes cell apoptosis of Wilms tumor cells

We found that Wilms tumor tissues had a much higher expression level of PBOV1 compared with adjacent normal tissues (**Fig 5A**). Similarly, we detected higher expression of PBOV1 in Wilms tumor cells (WiT49 and STA-WT3ab) compared with that in control HEK 293T cells (**Fig 5B**). To study the function of PBOV1, siRNA-targeting PBOV1 was used to suppress the expression of PBOV1 in WiT49 or STA-WT3ab cells. The knockdown efficiency was evaluated by western blot (**Fig 5C**). Functionally, we demonstrated that knockdown of PBOV1 suppressed cell proliferation and enhanced apoptosis in WiT49 or STA-WT3ab cells (**Fig 5D** and **5E**). While transwell assay revealed that inhibition of PBOV1 decreased the capability of cell migration and invasion in WiT49 or STA-WT3ab cells (**Fig 5F** and **5G**). Thus, the results suggested that PBOV1 acted as an oncogene in Wilms tumor.

**Figure 5.**
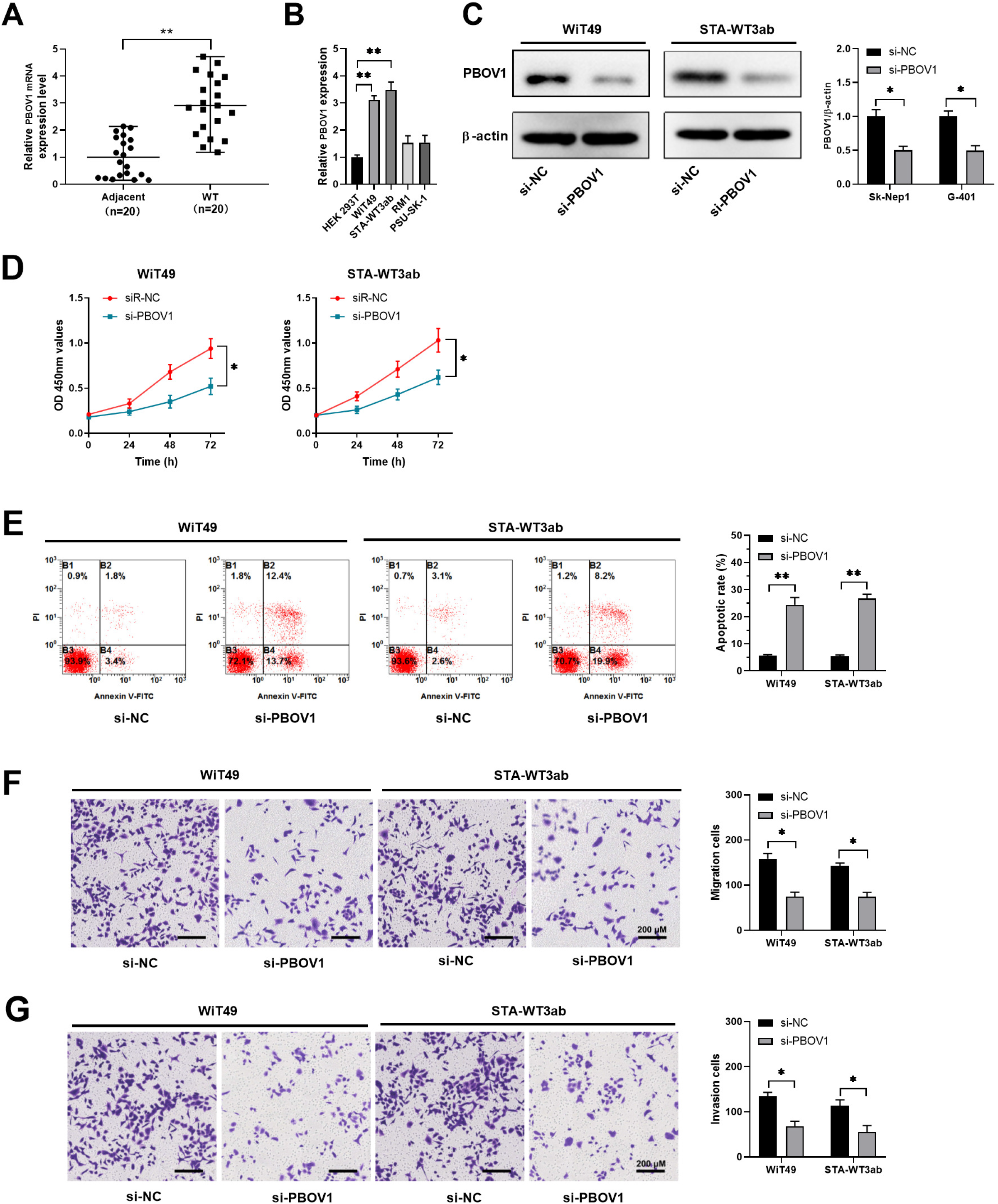
Knockdown of PBOV1 suppresses cell migration and invasion and promotes cell apoptosis of Wilms tumor cells. (A) The relative expression of PBOV1 mRNA in twenty pairs of Wilms tumor tissues and adjacent normal tissues was analyzed by qPCR. (B) The relative expression of PBOV1 mRNA in Wilms tumor cell lines (WiT49, STA-WT3ab, RM1, and PSU-SK-1) and control cell line HEK 293T was analyzed by qPCR. (C-G) WiT49 or STA-WT3ab cells were transfected with negative control (si-NC) or si-PBOV1. (C) The protein level of PBOV1 was analyzed by western blot 48 hours post-transfection. (D) Cell proliferation was analyzed by CCK8 assay. (E) Cell apoptosis was analyzed by Annexin V/PI staining and flow cytometry. (F, G) Cell migration and invasion were assessed by transwell assay. * *p* < 0.05, ** *p* < 0.01 vs. si-NC.

### Overexpression of PBOV1 antagonizes the tumor suppressor function of miR-27a-5p in Wilms tumor cells

To validate the functional relationship between miR-27a-5p and PBOV1, rescue experiments were performed. WiT49 or STA-WT3ab cells were transfected with miR-NC, miR-27a-5p mimic, or miR-27a-5p mimic+pcDNA-PBOV1. Whereas miR-27a-5p overexpression significantly decreased the expression of PBOV1, overexpression of PBOV1 together with miR-27a-5p rescued PBOV1 expression in Wilms tumor cells (**Fig 6A**). Functionally, miR-27a-5p mimic inhibited cell proliferation and PBOV1 overexpression abrogated the inhibitory effect or miR-27a-5p (**Fig 6B**). Conversely, miR-27a-5p enhanced cell apoptosis of WiT49 or STA-WT3ab cells. Overexpression of PBOV1 together with miR-27a-5p mimic showed reduced cell apoptosis and was comparable to that in cells transfected with miR-NC control (**Fig 6C**). In addition, overexpression of PBOV1 antagonized the inhibitory effect on cell migration and invasion of miR-27a-5p in WiT49 or STA-WT3ab cells (**Fig 6D** and **6E**). Taken together, the data suggested miR-27a-5p exerted its tumor-suppressive role in Wilms tumor cells via downregulating PBOV1.

**Figure 6.**
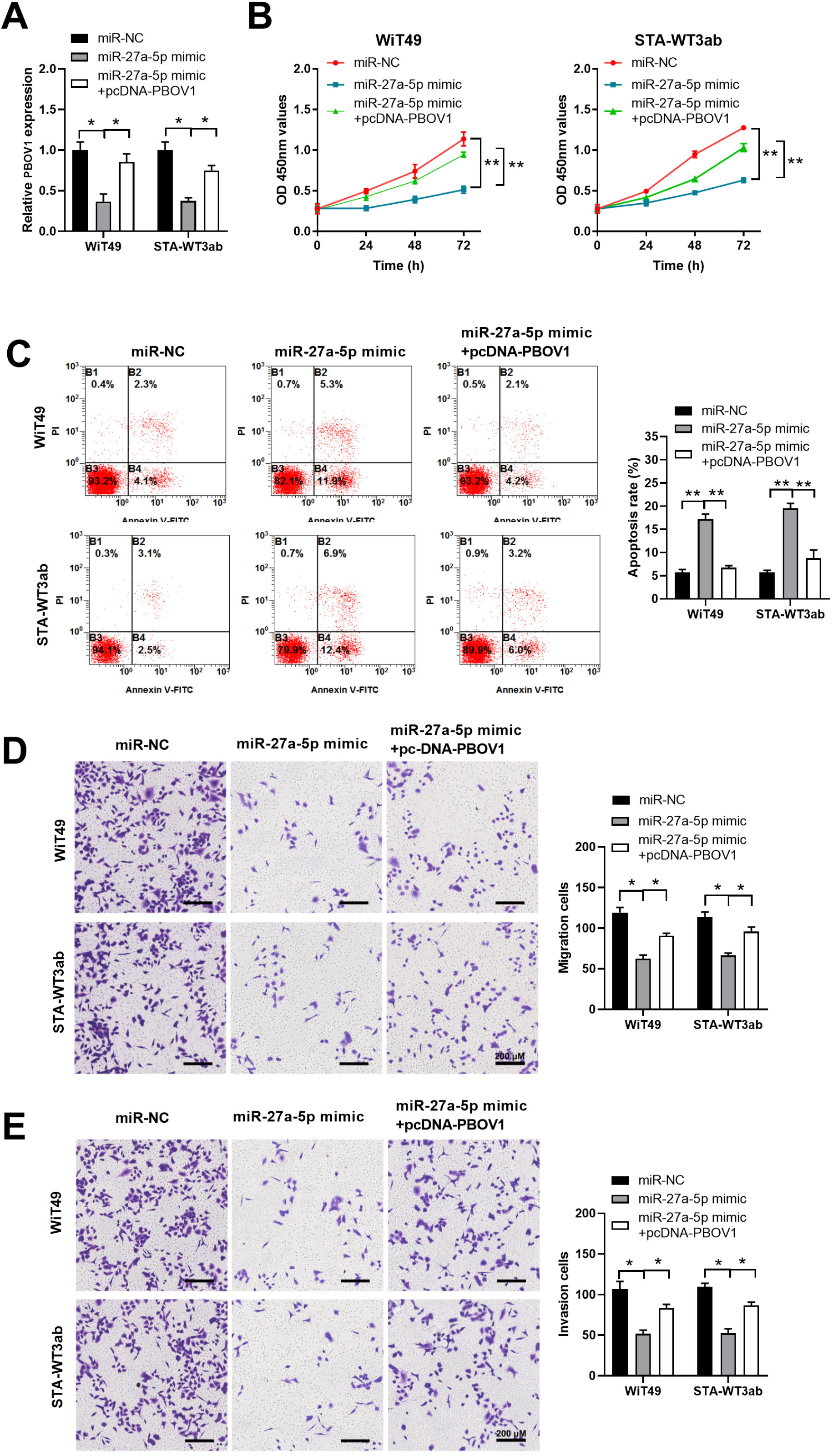
Overexpression of PBOV1 antagonizes the tumor suppressor effect of miR-27a-5p in Wilms tumor cells. WiT49 or STA-WT3ab cells were transfected with negative control (miR-NC), miR-27a-5p mimic, or miR-27a-5p mimic+pcDNA-PBOV1. (A) The relative mRNA level of PBOV1 in WiT49 or STA-WT3ab cells was analyzed by qPCR 48 hours post-transfection. (B) Cell proliferation was analyzed by CCK8 assay at indicated time points. (C) Cell apoptosis was analyzed by Annexin V/PI staining and flow cytometry. (D, E) Cell migration and invasion were assessed by transwell assay. * *p* < 0.05, ** *p* < 0.01.

## Discussion

Studies have shown that miRNAs play essential roles in tumorigenesis and metastasis(Si et al., 2019). MiRNAs also exerts the regulatory function in Wilms tumor and could be used as diagnostic markers and predictors for chemo-responsiveness(Schmitt et al., 2012; Watson et al., 2013). Here we reported that miR-27a-5p was low-expressed in Wilms tumor and miR-27a-5p overexpression suppressed Wilms tumor cell growth and metastasis. PBOV1 was demonstrated to be the direct target of miR-27a-5p and overexpression of PBOV1 abrogated the tumor-suppressive function of miR-27a-5p. Thus, our findings suggest a potential therapeutic target of miR-27a-5p in Wilms tumor patients.

MiR-27a-5p has been reported to suppress the tumorigenesis of multiple cancers including prostate cancer, small cell lung cancer, and cervical adenocarcinoma(Mizuno et al., 2017; Barros-Silva et al., 2018; Fang et al., 2018). Networks analysis showed that miR-27a-5p was dysregulated in Wilms tumor(He et al., 2016). In another study, Jenny A. Watson et al. showed that down-regulation of miR-27a was found in the high-risk Wilms tumors, which might be a predictor of chemo-responsiveness(Watson et al., 2013). We confirmed that miR-27a-5p was low-expressed in Wilms tumor. Consistent with the published data, the tumor-suppressive function of miR-27a-5p was validated both in vitro and in vivo. As overexpression of miR-27a-5p inhibited the growth and metastasis of Wilms tumor cells and promoted cell apoptosis.

Bioinformatics analysis predicted multiple potential targets of miR-27a-5p while we verified that miR-27a-5p mimics specifically inhibited the expression of PBOV1. PBOV1 was first identified as a human tumor-specific gene and associated with the clinical outcome of cancer patients(Krukovskaia et al., 2010; Samusik et al., 2013). The high expression level of PBOV1 promoted G1/S transition and enhanced cell proliferation in prostate cancer(Pan et al., 2016). The function of PBOV1 was also elucidated in hepatocellular carcinoma and overexpression of PBOV1 was correlated with poor prognosis of HCC patients, indicating PBOV1 as a prognostic biomarker of HCC(Xue et al., 2018). However, there are few reports regarding the regulation of PBOV1 in tumors. Zhang SY et al demonstrated PBOV1was regulated by miR-203 in fracture healing(Zhang et al., 2018). In rheumatoid arthritis, monocyte differentiation was controlled by lncRNA NTT/PBOV1 axis(Yang et al., 2018). In this study, we reported for the first time that PBOV1 was directly regulated by miR-27a-5p and PBOV1 overexpression antagonized the function of miR-27a-5p.

Though we demonstrated that miR-27a-5p/PBOV1 axis regulated Wilms tumor development and progression with both in vitro and in vivo evidence, there are several limitations in this study. First, it is of great importance to further study whether high expression of miR-27a-5p mediated the chemo-resistance in Wilms tumor or not. Second, whether there are other potential miRNAs involved in PBOV1 regulation remain unknown. Additionally, the signaling pathways involved in miR-27a-5p/PBOV1 axis in Wilms tumor need further investigation.

## Conclusion

MiR-27a-5p acts as a tumor suppressor via negatively regulating PBOV1 in Wilms cancer. Our data suggest that miR-27a-5p/PBOV1 might be utilized as a novel therapeutic target in Wilms tumor.

## Abbreviations

miRNA: microRNA;
3’-UTR: 3’-Untranslated region;
ATCC: American Type Culture Collection;
PI: Propidium iodide;
WT: Wilms tumor.

## Acknowledgments

**N/A**

## Authors’ contributions

Zheng-Tuan Guo and Qiang Yu conceived and designed these experiments. Chunlin Miao, Wenan Ge and Peng Li performed these experiments. Qiang Yu and Wenan Ge analyzed and interpreted the data. Chunlin Miao and Peng Li wrote the manuscript. Zheng-Tuan Guo and Qiang Yu revised the manuscript. All authors read and approved the final manuscript.

## Ethics approval and consent to participate

The ethics committee of the Second Affiliated Hospital of Xi’an Jiaotong University approved the study. The investigation conforms to the principles outlined in the Declaration of Helsinki and written informed consent was obtained from all participants.

## Availability of data and material

The datasets used and/or analyzed during the current study available from the corresponding author on reasonable request.

